# Neurophysiological Markers for the Acute Pain Model in Rats Under Urethane-induced Anesthesia

**DOI:** 10.1101/2025.08.27.672445

**Authors:** Dmitrii Andreevich Perevozniuk, Anna Yurievna Gorovaya, Kirill Sergeevich Smirnov, Lidia Mikhailovna Birioukova, Inna Stanislavovna Midzyanovskaya, Nikolay Vladimirovich Syrov, Igor Alexandrovich Lavrov, Mikhail Albertovich Lebedev

## Abstract

Certain neurophysiological mechanisms of pain can be investigated in anesthetized animals, and such a model is potentially useful for the development of pain treatment. Yet, the interference of pain-related neural patterns and anesthesia-related patterns should be understood. Here we studied the interplay between acute pain-induced and urethane-induced EEG activity. We analyzed the activity of the rats somatosensory cortex under urethane-induced anesthesia both before and after introducing a painful stimulus (formalin injection into the left hindlimb). Our analysis of the EEG parameters, such as amplitude, amplitude difference between the left and right hemispheres, and spectral entropy showed significant pain-related effects. The responses to painful stimuli depended on the anesthesia phase. These results are applicable to further research in animal models and to monitoring humans placed under general anesthesia.

## Introduction

The fundamental neurophysiological mechanisms underlying the ability to perceive and process painful stimuli have been extensively studied in both humans and animals. However, the complex interplay between pain mechanisms and the effects of anesthetics are still insufficiently understood. Understanding how the brain responds to painful stimuli while the organism is under anesthesia is critical to improving patient care, developing anesthesia and analgesia protocols, and expanding our knowledge of pain neurophysiology.

Urethane-induced anesthesia has been a valuable tool in animal research due to many advantages including its stability and long-lasting effects [2]. Unlike many other anesthetic agents, urethane does not exhibit a rapid onset and recovery, allowing for extended periods of stable anesthesia [2, 3]. Urethane cannot be used in humans though due to its high oncogenic and other side effects [4]. The mechanisms by which urethane affects cortical activity, as evidenced by alterations in electroencephalographic (EEG) activity, and the nature of its effects on pain processing remain poorly understood.

Previous studies have shown that brain activity during urethane anesthesia is characterized by two distinct phases of EEG activity [5]. The initial phase, known as the desynchronization or activation, is characterized by a presence of high-frequency oscillations, resembling those observed in awake animals [5]. The second phase, referred to as synchronization or suppression, is manifested by the presence of low-frequency, high-amplitude oscillations. Examples of these phases in EEG are illustrated in Figure 1. The resemblance of these EEG patterns to normal sleep cycles led scientists to suggest that the urethane-induced anesthesia could be used as a model of sleep [6]. However, recent studies have questioned the validity of this model [7] and the alternating EEG phases under urethane anesthesia remain a matter of debate. Specific interest in disentangling these mechanisms is fostered by the analgesic effects of urethane. Previous research has demonstrated that various anesthetic agents can modulate pain perception and alter EEG patterns [8] but the specific effects of urethane on pain-induced EEG activity have not been thoroughly investigated.

**Figure 1.**
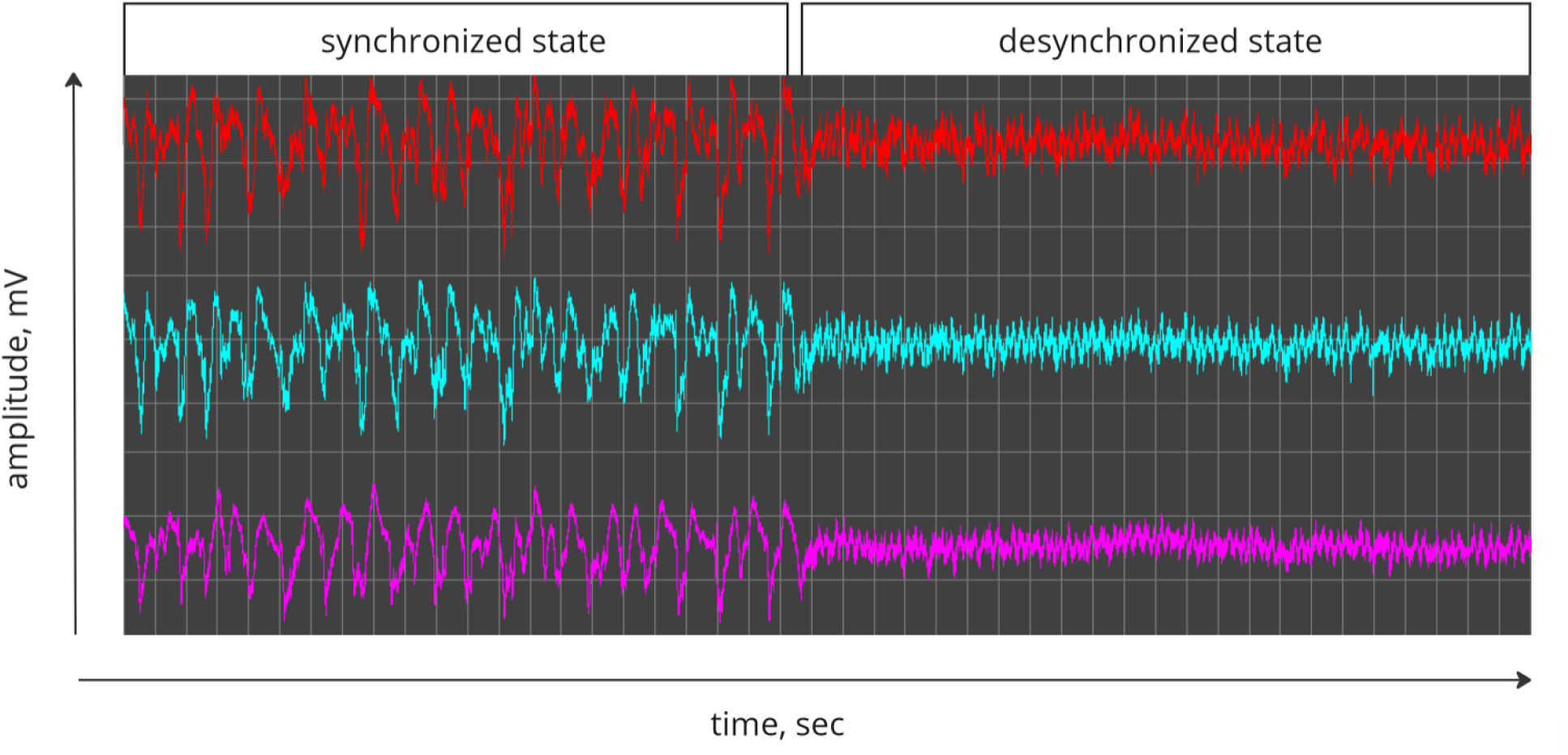
Example of synchronized and desynchronized states under urethane anesthesia. Three EEG channels to the rat cortical areas: right and somatosensory cortex, and right occipital cortex (top to bottom). EEG recordings were conducted with epidural electrodes.

**Figure 2.**
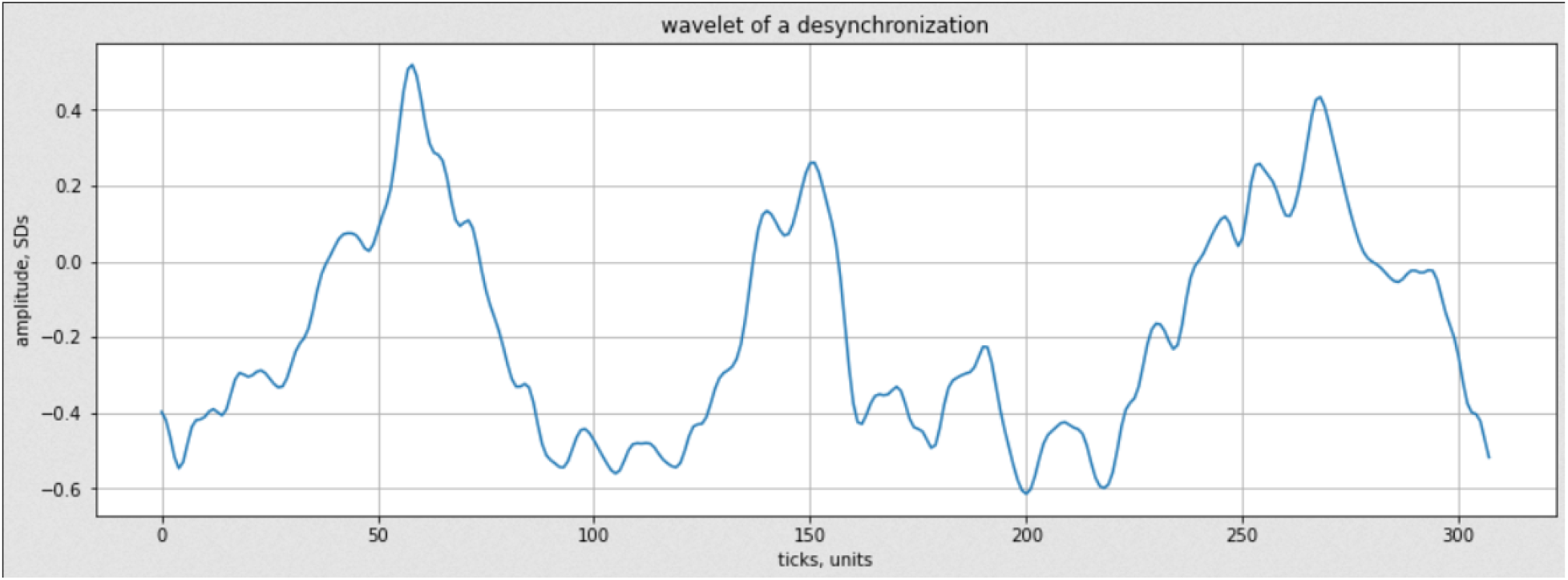
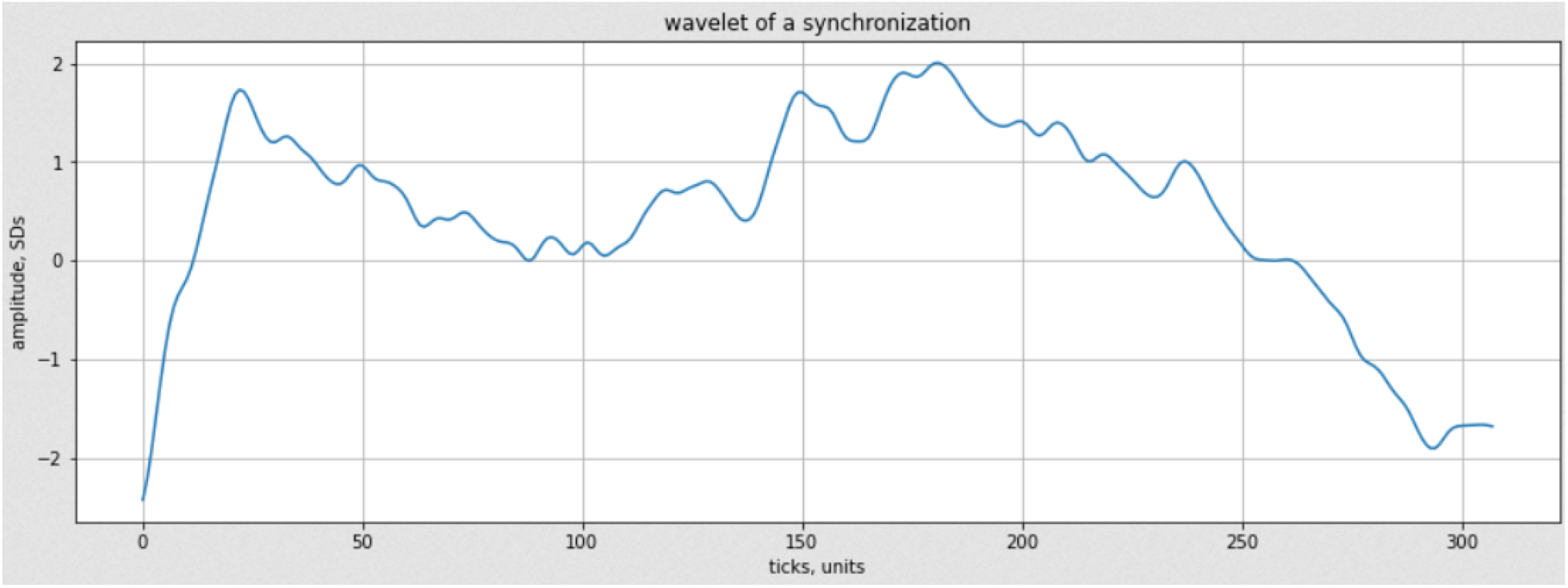
Wavelets used for convolution: first graph – wavelet taken from a desynchronization phase, second graph – wavelet taken from synchronization phase. Both wavelets were smoothed using two-way exponential smoothing with alpha = 0.2.

The current study addresses this knowledge gap by investigating the changes in EEG oscillatory patterns in response to painful stimuli under urethane anesthesia. By examining brain activity recorded from rats’ somatosensory cortices before and after the introduction of a painful stimulus, we aimed to better understand the interplay between pain-induced and urethane-induced cortical activity. To induce acute pain, we employed the formalin injection model, which is well-established for studying acute pain in animals [9]. This model produces a biphasic pain response, with an initial acute phase followed by a longer-lasting inhibitory phase [9]. Focusing on the first, which lasts approximately from one to two hours, we aimed to uncover immediate effects of pain on EEG activity of the rat somatosensory cortex and hemispheric specificity of these effects. By analyzing various EEG oscillations amplitude in both activation and deactivation phases, its interhemispheric differences, and spectral entropy as value reflecting complexity of signal, we aimed to identify specific neurophysiological markers of pain under an anesthesia.

## Methods

### Animals

This study was performed in 13 adult Wistar male rats. The rats were bred and housed in the vivarium of the Institute of Higher Nervous Activity and Neurophysiology of RAS. The experimental procedures conducted in this study were executed in compliance with the regulations stipulated by EU Directive 2010/63/EU for animal experimentation [1]. The animal ethics committee of IHNA approved the protocol (No. 4) on the 26th of October, 2021. Throughout the study, the rats were maintained under the standard conditions adhering to a 12:12 hours light-dark cycle with lights turned on at 08:00. Adequate ventilation and airing systems were integral parts of their housing environment. Prior to the surgical procedures, the rats were kept in small same-sex groups with 3-4 rats per cage, and they had unrestricted access to food and water.

### Surgery

In order to record electrical brain activity, epidural electrodes were chronically implanted. The surgery was performed under general isoflurane anesthesia in a Kopf stereotaxic apparatus. Three active epidural screw electrodes were implanted over the left/right somatosensory cortex (AP 0; L 3) and occipital cortex (AP −6; L 3). The reference screw electrode was placed above the cerebellum. After surgery, animals were kept individually on a 12/12 light/dark cycle with unrestricted access to food and water. The recovery period lasted 14 days.

### EEG Recording

All recording sessions were performed in the noise-isolated room. Rats were placed in Plexiglas cages (25 cm × 60 cm × 60 cm). Urethane anesthesia was induced (1.2 mg/kg, 10 ml per 1 kg of body weight). The signals were acquired with a multi-channel amplifier (PowerLab 8/35, ADInstruments, Sydney, Australia) via a swivel connector. The signals were digitized at 2000 samples/s rate on each channel. After 45 mins from injection, EEG recordings started. Firstly, 2 hours of baseline activity was recorded. Secondly, a pain stimulus was administered by injecting a 5% formalin into the left hindlimb. Following that, 1 hour of pain induced activity was recorded. At the end, all rats were euthanized using DCL concentration of chloral hydrate.

### Data processing

Manual markup of the data was performed in EDF browser software (https://www.teuniz.net/edfbrowser/). All other stages of data-processing were performed using Python programming language and such libraries as MNE-python, NumPy, Pandas, SciPy, math, matplotlib, sklearn, pingouin, seaborn.

Since phases of urethane-induced anesthesia heavily affect EEG traits (image 1]), we distinguiahed synchronization and desynchronization phases. We were interested in formalization as well as automatization of this task to get rid of variance introduced by the human factor. For this, we developed a random forest type classifier [10]. This algorithm required training and testing subsets. Initially, out of N recordings, 6 were marked up manually, thus 4 were used as training and 2 as testing.

The preprocessing stage consisted of several steps. First, a band-path filtration in the range 0.5-100 Hz was applied. Second, recordings were standardized – by dividing signals’ amplitude values by standard deviation of this channel, values were converted from mV to SDs. These steps allow for a comparison between recordings with different signal quality, as well as negating some artifacts.

Before passing into the classifier, EEG recordings were processed in the following way: for every 1-second-long epoch, spectral entropy value was calculated, along with two distinct wavelet values. Every epoch overlapped with one before and one after, so that the first 20% of epoch’s values are the same as the last 20% of previous ones.

Spectral entropy was computed using equation 1. Individual spectra were generated for each channel, and the resulting entropy values were averaged across all three channels. This metric serves to quantify how power spectral density is distributed within a signal, indicating the level of information or complexity present in its frequency domain. By analyzing the distribution of power across different frequencies within the signal, spectral entropy offers insight into the signal’s spectral makeup. While initially developed for signal analysis in cryptography and cryptanalysis, spectral entropy has also demonstrated utility in EEG analysis. The spectral composition of EEG signals varies significantly based on the individual’s physiological state, making metrics that characterize these properties valuable to physiologists and clinicians alike. Higher entropy values typically imply a spectral resemblance to white noise, suggesting a lower prominence of the slow-wave band compared to higher frequency bands. This phenomenon is associated with a shift in the inhibition/excitation balance towards excitation. In essence, entropy serves as a general metric linked to the balance between inhibition and excitation [11].

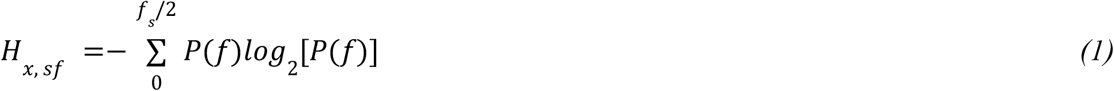

Equation 1. Spectral entropy calculation.

Where Hx, sf is entropy value for interval x with sample rate of sf. P(f) - normalized PSD.

As for the wavelet values, we used two distinct wavelets – one was a smoothed 1-s interval from the synchronization phase and the other was a similar interval from the desynchronization (image 2). Smoothing was performed using a custom script for two-way exponential smoothing, according to equation 2. That was a two-way smoothing applied in both directions to negate a lag introduced by one-way smoothing. Those wavelets were convolved with an epoch to get values describing similarity between a signal interval and a wavelet.

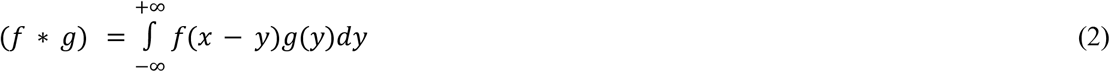

Equation 2. Convolution.

Where *f* and *g* are functions that are to be convolved, and x and y are independent variables.

All of the values described were calculated simultaneously for each of three channels and an epoch was described as a median of each value along three channels.

These values were normalized using a standard scaler from sklearn library, so that the median of each variable was equal to 0 and standard deviation was equal to 1.

Altogether, the classifier was presented with a dataset where three values which describe an epoch were aligned to binary markup of phases (for a training/testing subsets). Binary markup consisted of 1’s, which corresponded to the epochs where desynchronization was observed and 0’s, with a synchronization in the corresponding time window.

We intentionally did not include amplitude of the signal into the classifier input. Since further steps will include it as a dependent variable under different conditions, using amplitude of a signal to distinguish phases could have introduced some sort of a bias.

The classifier was trained to predict whether a desynchronization (state ‘1’) or synchronization (state ‘0’) was observed in a particular epoch, using three of the described above values. We chose random forest classifier model from sklearn.ensemble. Classifier was trained on 4 manually marked recordings, using RandomizedSearchCV optimizer [12] to find optimal state. Accuracy scoring was used. Optimal model for our task performed with accuracy of 0.96, f1 score of 0.94, AUC score of 0.99, 0.95 recall and precision of 0.99. All of the parameters listed above were calculated for testing subset (2 recordings). The model assigned approximately equal weights to all three input values (0.31 for entropy, 0.38 for desynchronization wavelet and 0.3 for synchronization wavelet), thus proving that input parameters necessary for effective discrimination were selected correctly. After achieving said values, the model was retrained using all 6 manually marked recordings.

Additionally, classification results were adjusted using two thresholds, one for minimal length of desynchronization and the other for minimal synchronization length. Both were set to 10 s, since both phases tended to last much longer [8, 9, 10]. The markup in those intervals which did not surpass one of the thresholds was changed to the opposite. For example, if the interval of desynchronization lasted less than 10 s, 1’s indicating desynchronization in time windows corresponding to said intervals were flipped to 0’s (and vice versa). This filtration effectively merges two nearby intervals if the interval of opposite type of activity separating them lasts less than 10 s. Such filtration allowed us to get rid of small intervals of false-positives and false-negatives (example on images below), increasing accuracy up to 0.98. Example of said filters’ result present in Figure 3.

**Figure 3.**
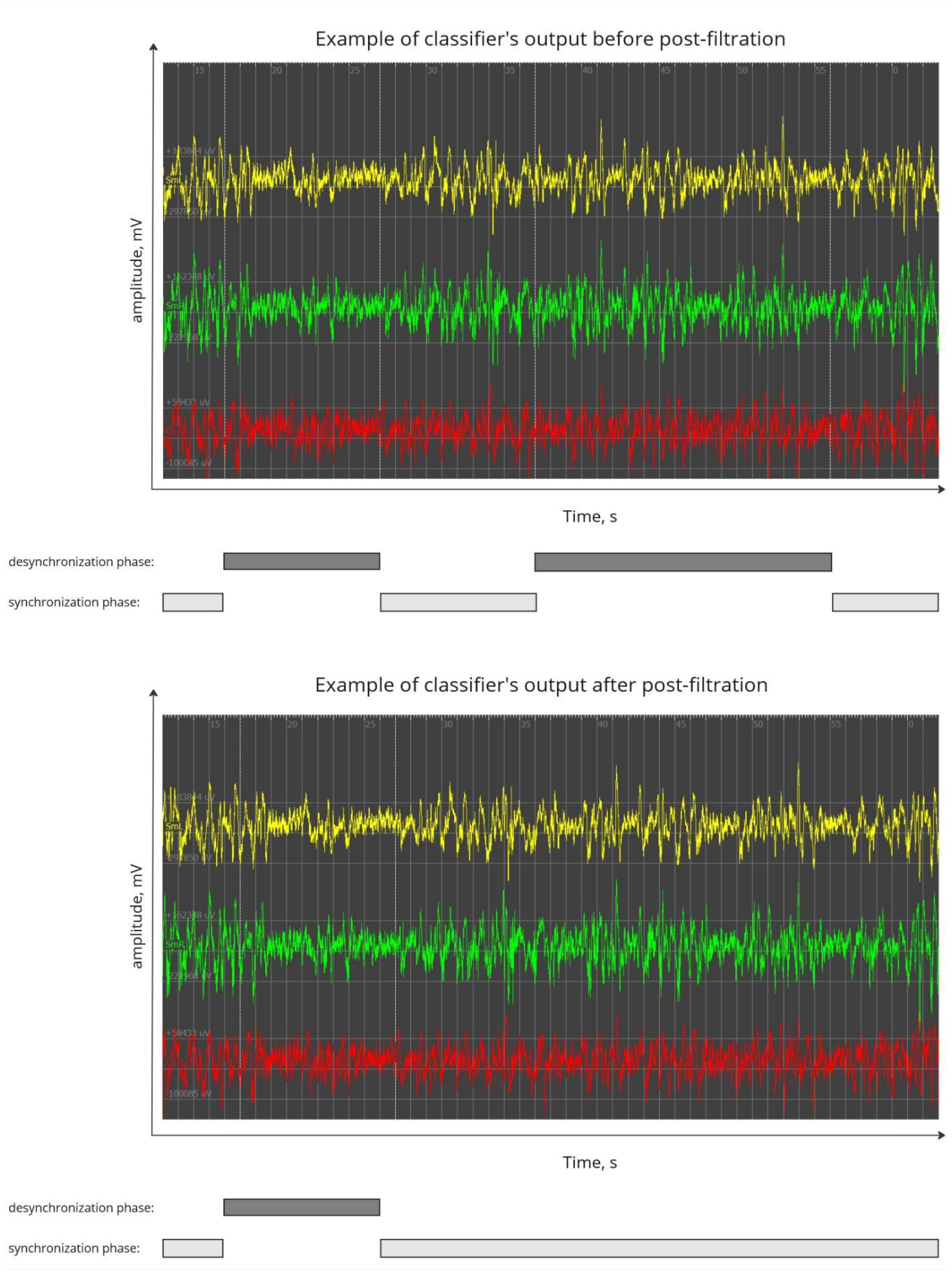
Illustration of post-filtration results in negating short intervals of false-negatives and false-positives.

Using the described approach, a total of 24 recordings were marked, each of which lasted for 2 hours.

### Data acquisition

For further analysis, each marked recording was sampled according to following algorithm:

1. 1000 epochs lasting for 20 s were sampled from each recording, 500 epochs from each phase.
2. Epochs were sampled in such a fashion that they cover each phase evenly and spacing between them is always equal or more than 10 seconds. Epochs are located at least 10 seconds from the beginning or the end of the phase.
3. For each epoch we calculate and store a list of parameters – median power in low (<20Hz) and high (>20Hz) power bands for both hemispheres, cross-correlation between hemispheres, spectral entropy for each hemisphere.
4. Depending on whether the EEG was recorded in baseline or after formalin injection, corresponding value is also put into the dataset, as well as ID of a rat from which EEG was recorded.

As the result, we acquired a data frame, in which each 20-s epoch was characterized by cross-correlation and power in different frequency bands and different hemispheres, coupled by ID of an animal, mark of whether a pain was induced and from which phase this epoch was sampled.

The overall schematic representation of the experimental pipeline is depicted in Figure 4.

**Figure 4.**
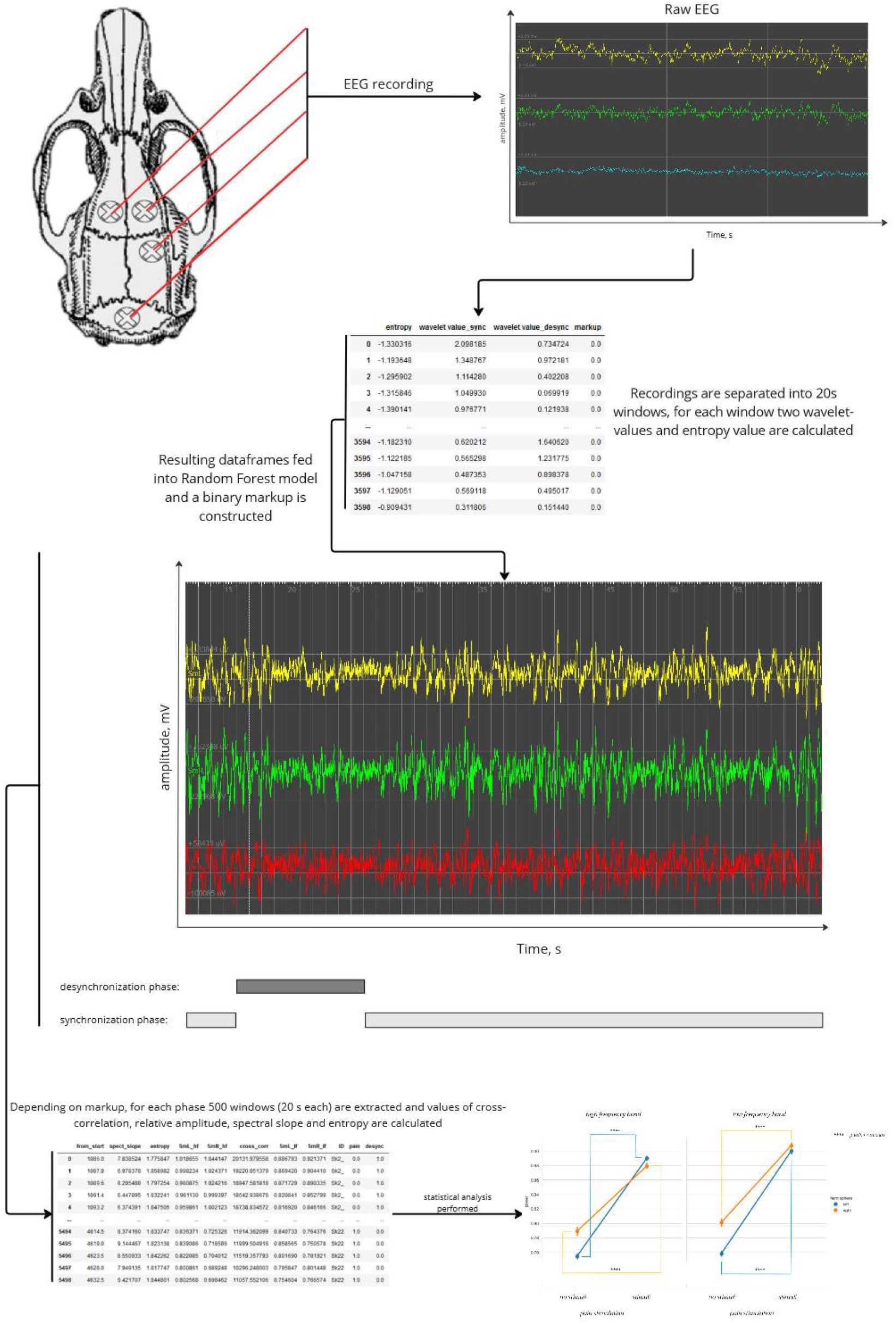
Experimental pipeline.

## Results

### Phase intervals lengths dynamics

Firstly, we analyzed the dynamics of synchronization and desynchronization intervals. For recordings of different length to be comparable, length of each interval was divided by total length of particular recording. We observed statistically significant differences (p-value<<0.0001, Mann-Whitney U-test, Bonferronni correction) between baseline and pain-induced states (Figure 5). Significant difference (p-value<0.05, Mann-Whitney U-test) was also shown for numbers of intervals per recording (graph 2). Both those results together show significant fragmentation of phases and/or more often switching between those while pain stimulus is administered.

**Figure 5.**
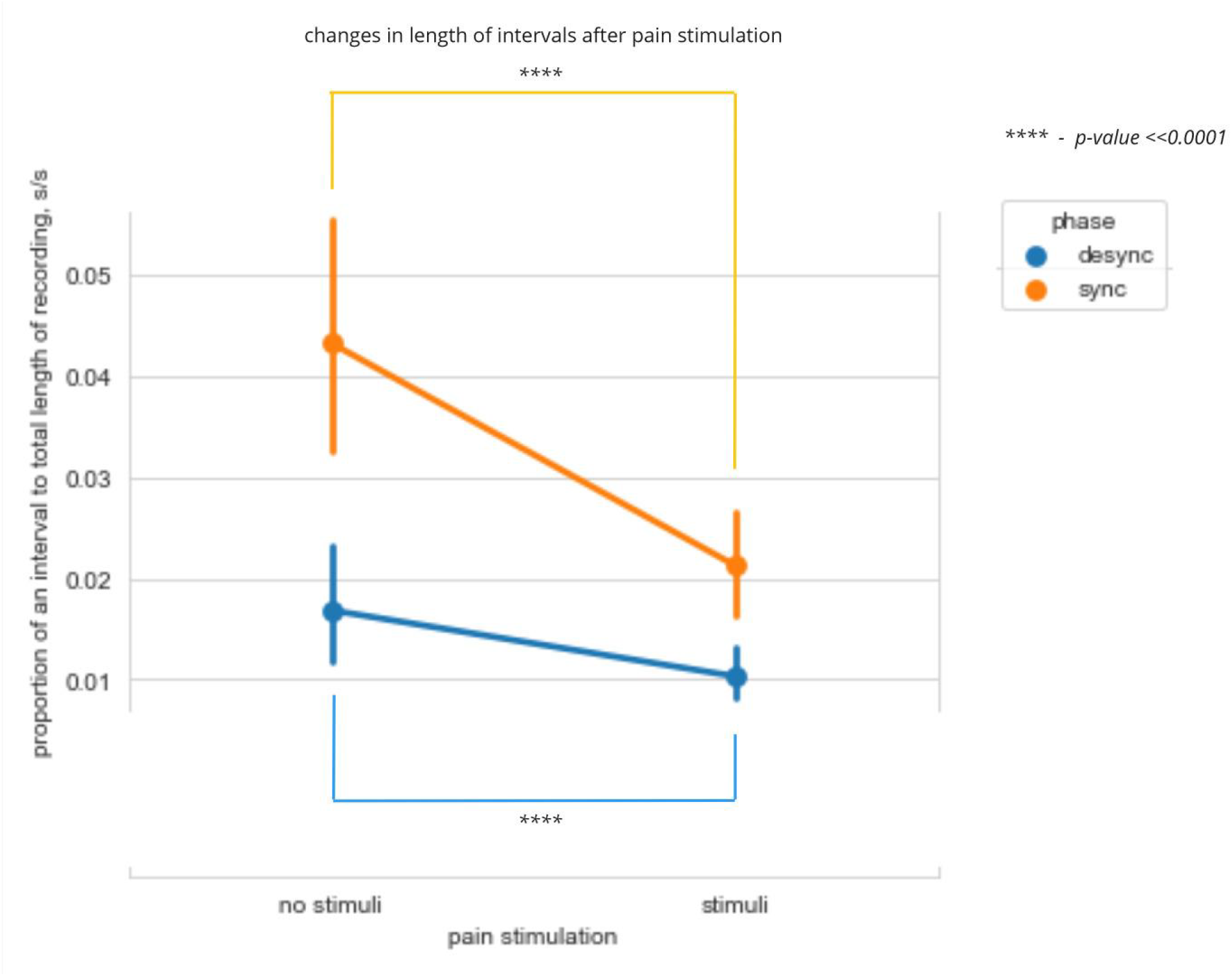
Differences in total length of synchronization and desynchronization intervals between baseline and pain-affected animals. Length of intervals (axis Y) is given in proportion to total length of particular recording (s/s). Statistical significance is reached for Mann-Whitney U-test, corrected for the number of comparisons (2) using Bonferroni correction.

**Figure 6.**
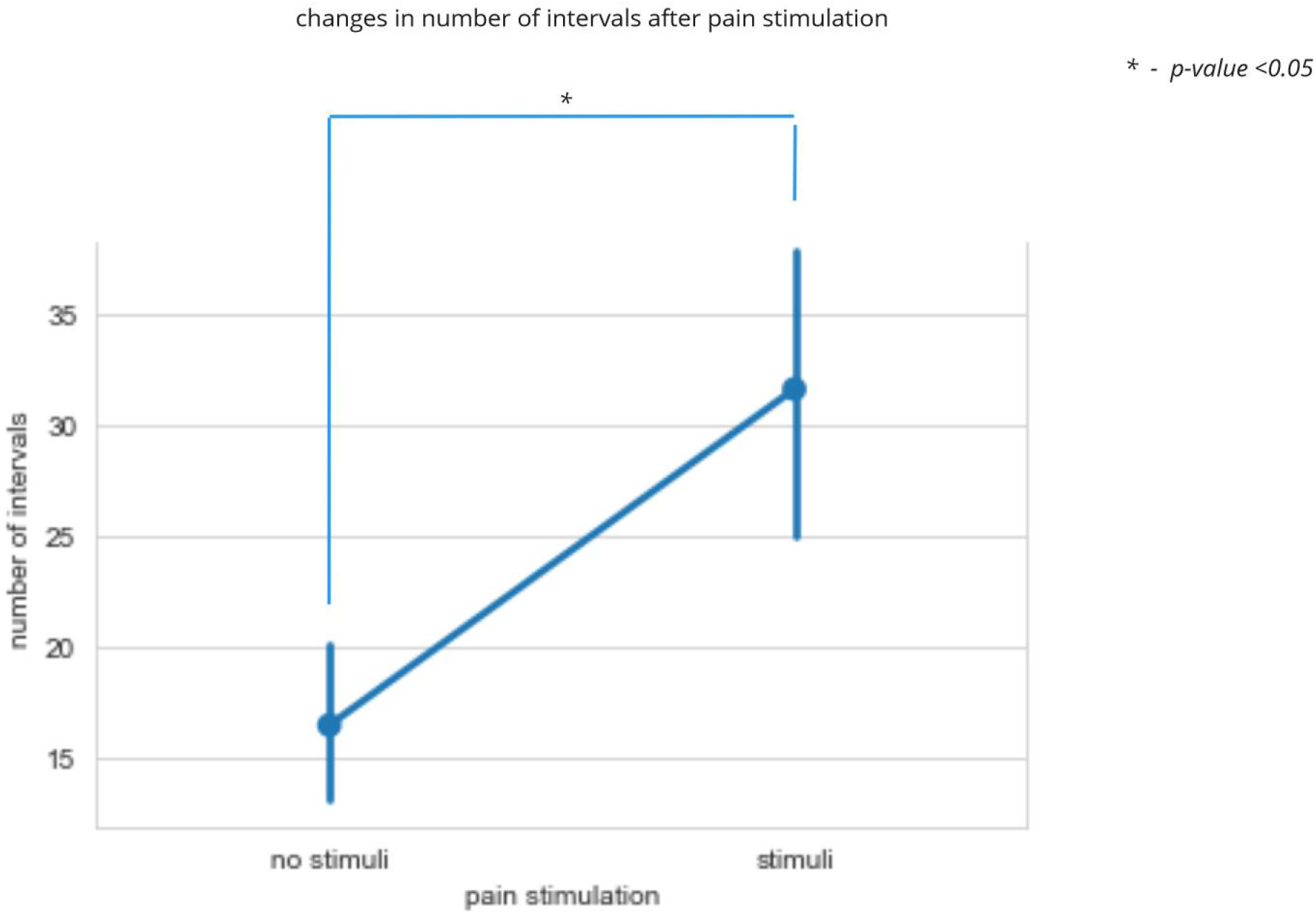
Differences in the number of synchronization and desynchronization intervals between baseline and pain-affected animals. Statistical significance is reached for Mann-Whitney U-test.

To test whether pain stimulation changes proportion of phases, we performed Fisher exact test and received significance (p-value<<0.0001, Fisher exact test, no correction). Therefore, resulting change of proportion from synchronization being 75,54% (*CI*_95_[49.07%:85.15%]) of recording to 72.79% (*CI*_95_[48.63%:79.88%]) of recording is to be considered further.

### EEG properties

To assess the effects of different nature, we analyzed the resulting dataset with ANOVA, including amplitude of a signal as dependent variable and hemisphere, phase of narcosis, pain stimulation and frequency bands as predictor qualitative variables. Since 14 comparisons take place in this test, all p-values (listed in p-unc column) were multiplied by the number of comparisons, thus adjusting for multiple comparisons according to Bonferroni post-hoc correction. While significance was achieved in a multitude of the interactions listed below (p-value<0.0001, Bonferroni correction), higher-order interactions should be analyzed and interpreted first. Since significant high-order interaction means that parameters affect each other and should be interpreted together. In our case, highest-order interaction between pain, narcosis phase, frequency band and hemisphere is statistically insignificant. This means that there is no such interaction and lower-level dependencies should be analyzed. Similarly, interactions between frequency band, pain and hemisphere and phase, frequency band and hemisphere are all insignificant. On the contrary, interaction of factors of pain, anesthesia phase and hemisphere are of importance (p-value < 0.0001, Bonferroni correction). This result can be interpreted so that amplitudes of a signal in different narcosis phases in different hemispheres are affected by pain in different fashion (we will delve deeper into this particular interaction in next paragraph). Similarly, statistical significance was achieved for the interaction between pain stimuli, narcosis phase and frequency band. Meaning difference in amplitude of a signal in response to a pain in different power bands and in different narcosis phases.

Results of individual factors should be analyzed separately. Although all four of the predictor variables show significance, we should also estimate the eta-squared value, representing the amount of variability explained by a factor. Indeed, the variable which explains highest variability is pain stimulus (eta-squared = 0.109, p-value<0.0001, Bonferroni correction).

For better visualization, we performed pairwise comparisons with amplitude of a signal as dependent variable while grouping by all four of predictors (Figures 7, 8). Significance (p-value <<0.0001, t-test, Bonferroni correction) was achieved when comparing the same hemispheres between intact and pain-stimulated for both phases of a narcosis (separated by frequency band, too). Interestingly, we observed a significant difference between hemispheres while intact (p-value <<0.0001, t-test, Bonferroni correction) that was gone after introduction of the stimulus. After addressing this topic with the surgeon who performed electrode implantation, we came to the conclusion that this anomaly (which persists in all rats and recordings) can be explained by positioning of the ground electrode. While both hemispherical ones were located symmetrically, that one was moved slightly to the right side to avoid massive blood loss from sinus suprasagitalis. If corrected for this effect, amplitude from left hemisphere is initially insignificant compared to right one (p-value>0.05, t-test, Bonferroni correction) and reaches significantly higher amplitude after pain stimulation (p-value <<0.0001, t-test, Bonferroni correction).

**Figure 7.**
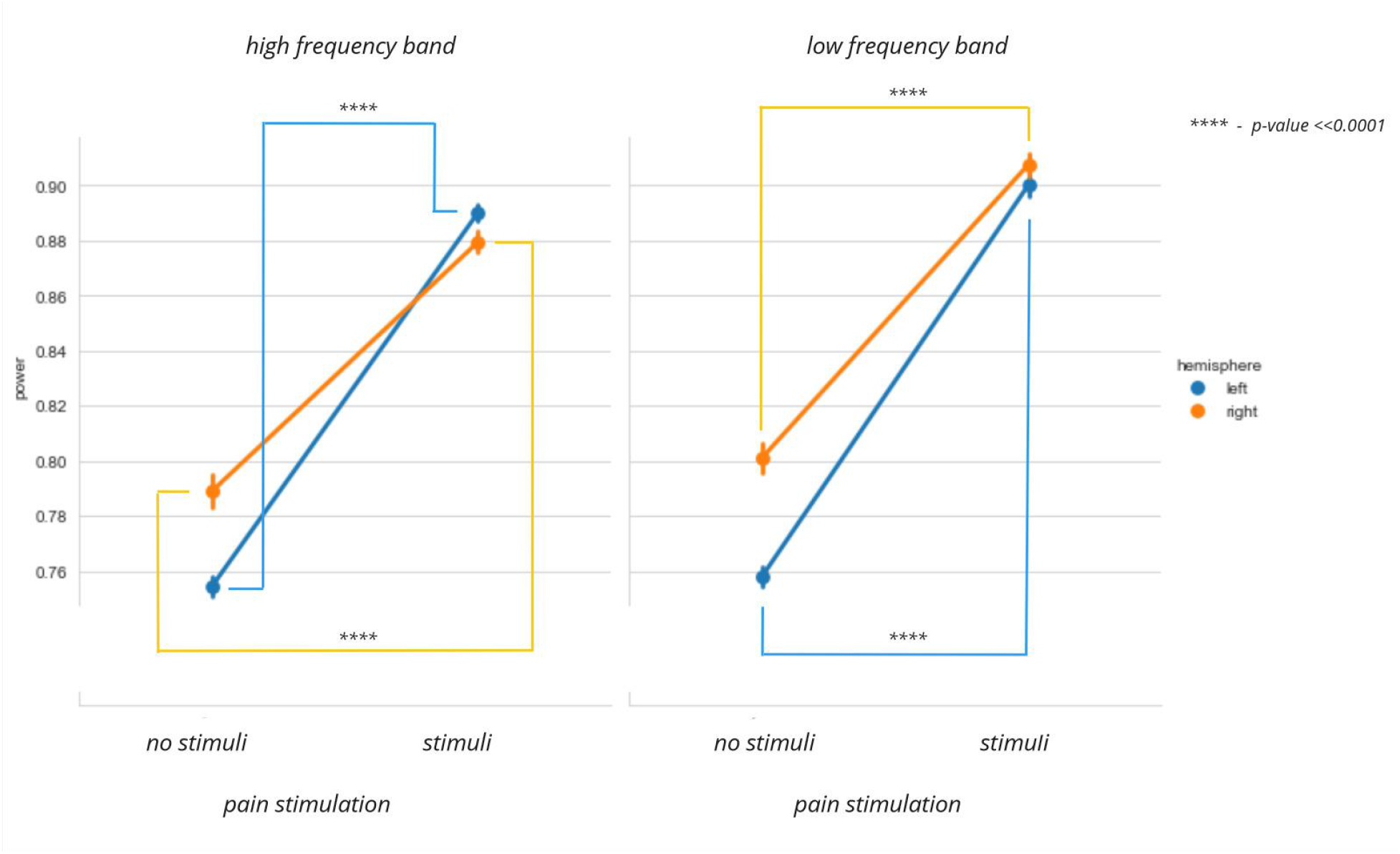
Representation of signal amplitude in different hemispheres and bands during the synchronization phase. (High frequency > 20 Hz power-band, low frequency 0.5-20 Hz power-band). Statistical significance is reached for t-test, corrected for the number of comparisons (8) using Bonferroni correction.

**Figure 8.**
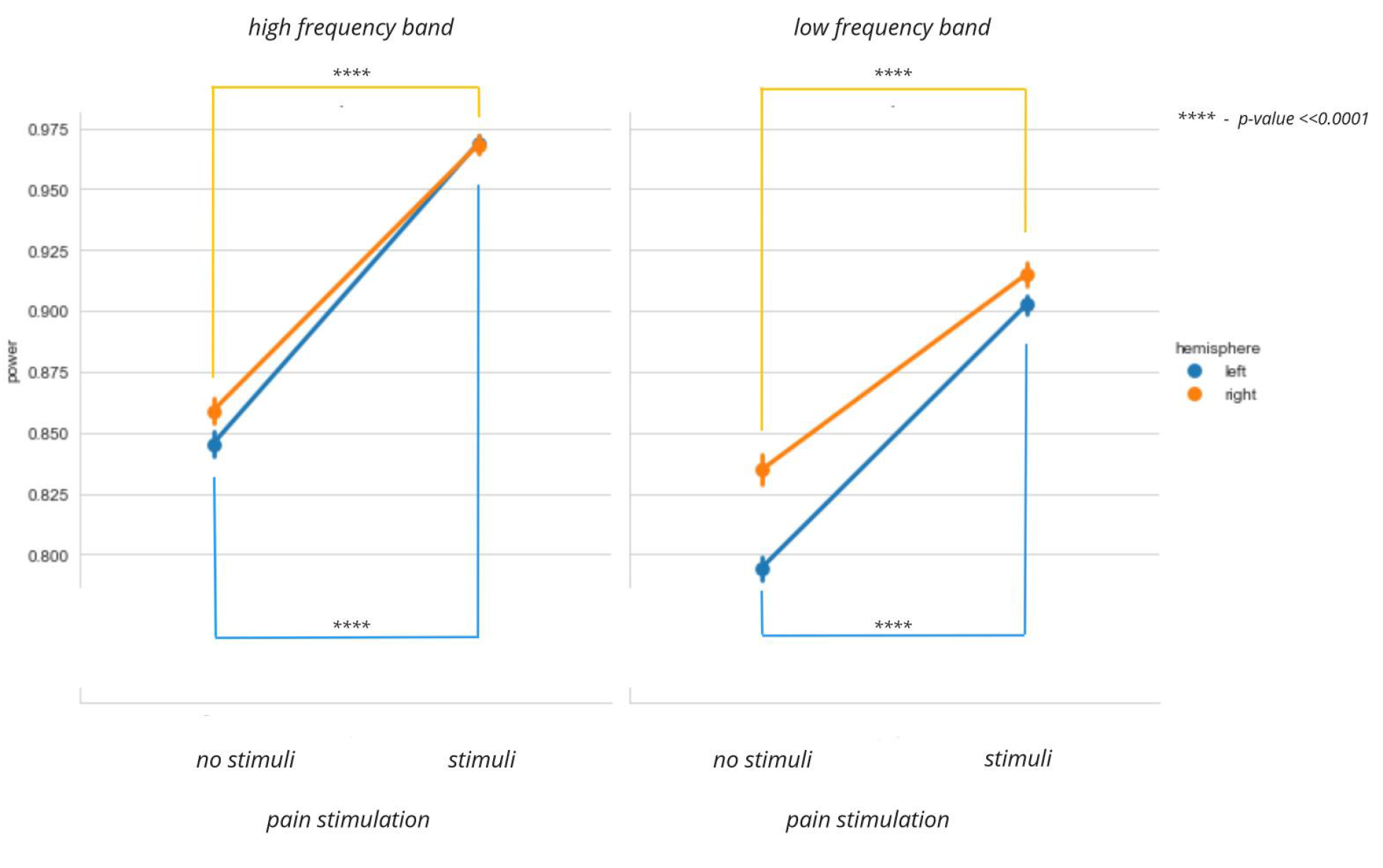
Representation of signal amplitude in different hemispheres and bands during the desynchronization phase. Statistical significance is reached for t-test, corrected for the number of comparisons (8) using Bonferroni correction.

**Figure 9.**
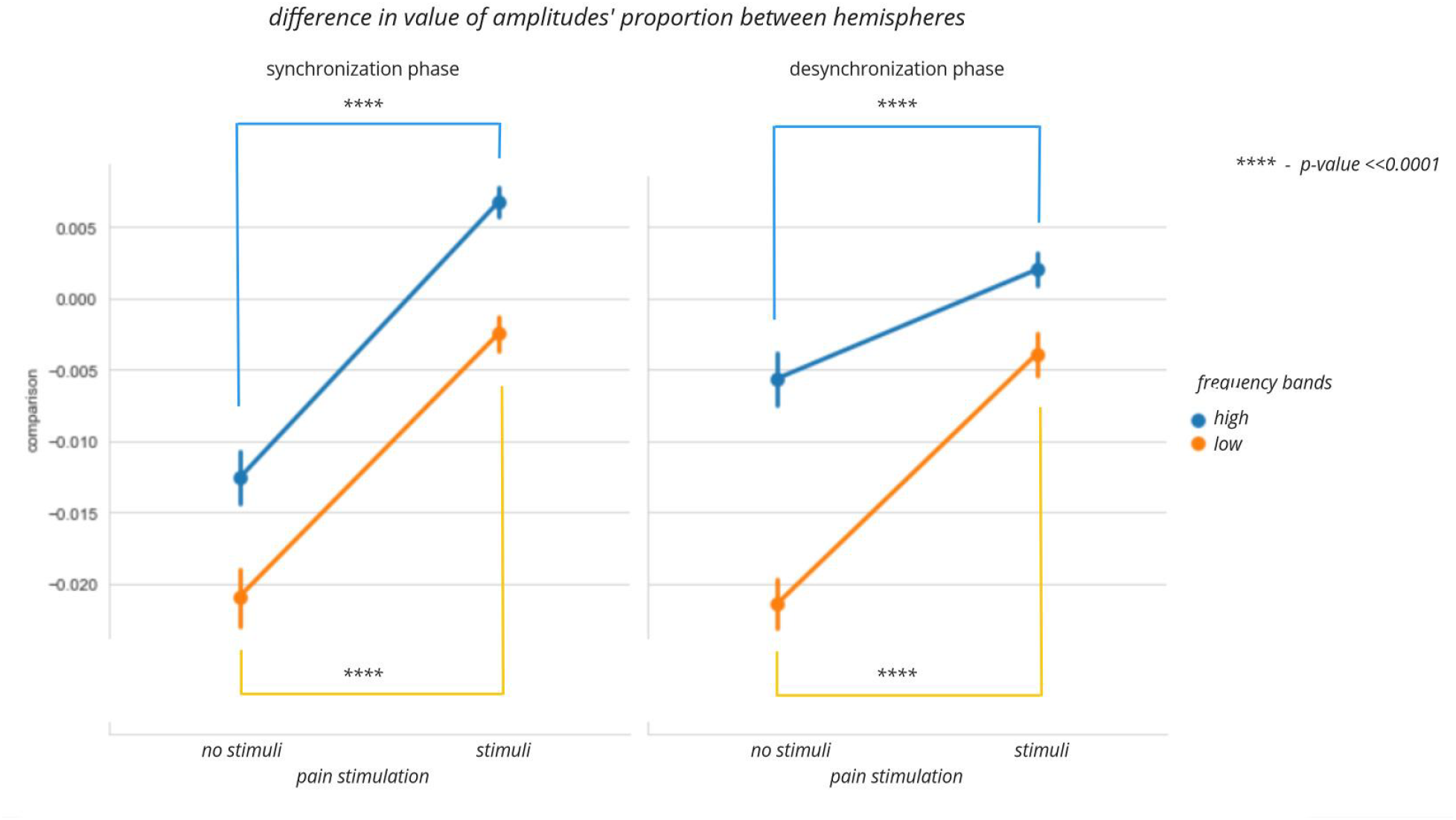
Graph representing differences in comparison value: left graph represents synchronization, while right graph depicts desynchronization state. (Note that, since ‘comparison’ value is calculated as difference between left and right hemispheres’ amplitudes divided by sum of those, its growth displays either disproportional decrease of right hemisphere’s activity or disproportional increase of left-sided amplitude. Possibly even both). Statistical significance is reached for t-test, corrected for the number of comparisons (4) using Bonferroni correction.

**Figure 10.**
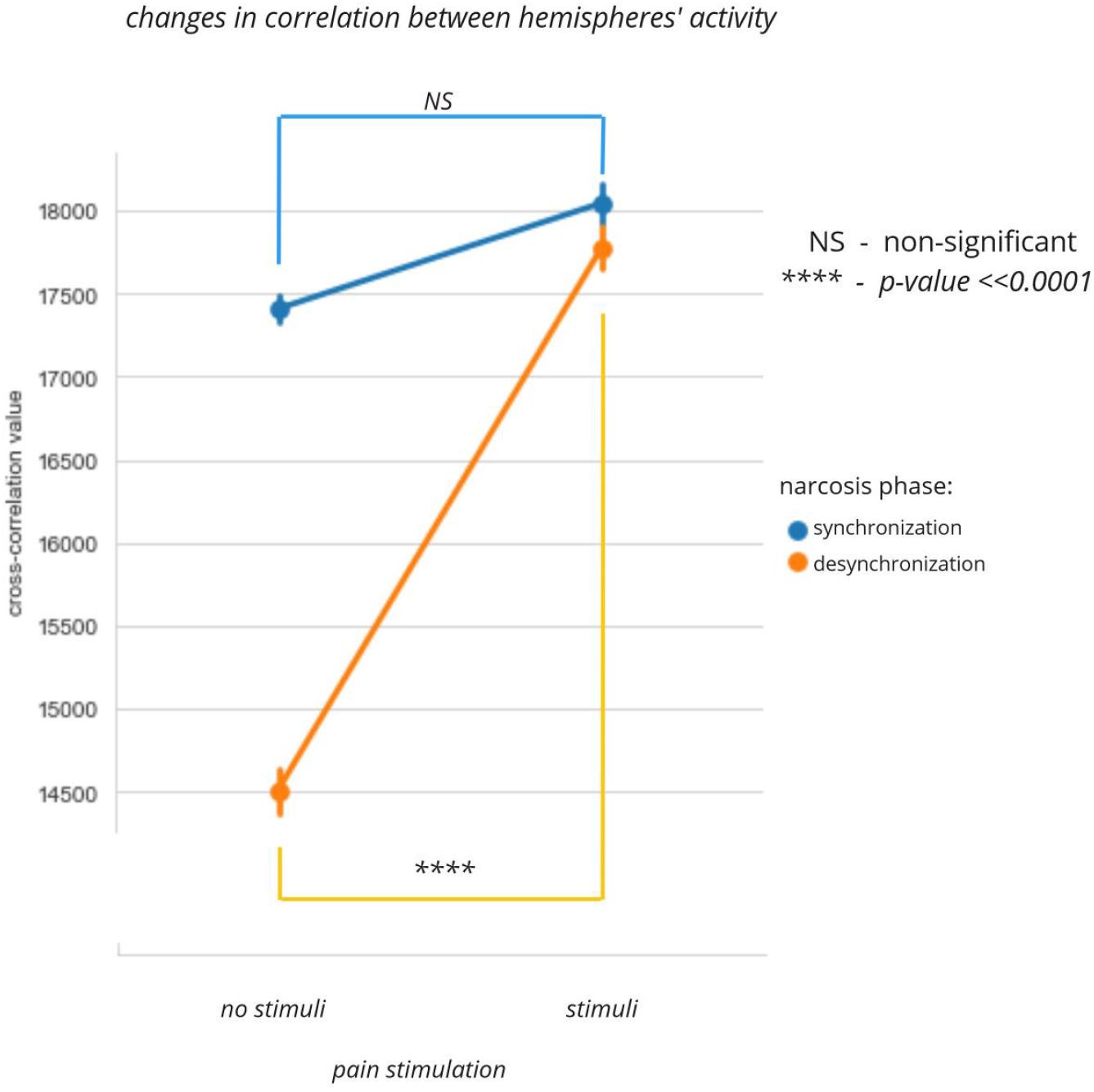
Graph depicts changes in cross-correlation value after pain stimulation during desynchronization and synchronization phases. Statistical significance is reached for t-test, corrected for the number of comparisons (2) using Bonferroni correction.

After that, we introduced a new dependent variable to further investigate different hemispheres’ responses. For that we calculated a ‘comparison’ value, which was equal to power in the left hemisphere minus power in the right hemisphere and the result divided by the sum of those two. This value was calculated for each interval for both frequency bands separately. Since it describes the power of hemispheres relative to each other, we expect it to grasp potential pain-related asymmetrical responses better.

Performing ANOVA once again, we observe statistical significance (p-value<0.0001, Bonferroni correction) for the highest order interaction, limiting our ability to interpret other interactions of factors. Interestingly, in previous analysis akin interaction between all four predictors did not show any significance. Looking at single factors, we observe the lack of significance for the phase of narcosis. ALthough quite unexpected at first, this observation aligns well with the fact that anesthesia is a generalized impact. It indeed greatly affects spectral properties of an EEG, as well as introducing phase switching. But those effects are not unique for a certain hemisphere and thus they do not affect comparison of hemispheres’ activities (while showing effect on raw amplitude).

Going further, we once again observe the highest eta-square for pain stimulation, confirming the ability of introduced comparison value to grasp response to pain stimulation.

Following that, pairwise comparisons using t-test were performed (results on graph 5). Significant growth of comparison value (p-value <<0.0001, t-test, Bonferroni correction) was observed, meaning that hemispheres react differently and the hemisphere which is contralateral to stimulation (right one) shows lower amplitude in response to it. Once again the left hemisphere seems to show lower baseline amplitude of a signal, registered as lower comparison value. As described above, this effect comes from positioning of ground electrodes and thus appears in both raw amplitude and comparison value levels.

Our dataset also held a cross-correlation value for each one of intervals. We performed ANOVA with it as a dependent variable and narcosis phase and pain stimulation as independent ones. Statistical significance was achieved both for interaction and individual factors alike (p-value<<0.0001, Bonferroni correction). These results suggest different responses to pain stimulation in different phases of narcosis in regard to cross-correlation. Further, pairwise comparisons were performed using t-test. After correction for multiple comparison, we no longer observe significant changes in cross-correlation during the synchronization phase, as opposed to desynchronization (graph 6). While high correlation is expected during slow-wave activity, lack of any significant reaction is remarkable. On the other hand, during the desynchronization phase, cross-correlation seems to react very vividly.

## Discussion

Our investigation into the different cortical responses observed during the distinct phases of urethane anesthesia clarify the dynamic nature of pain processing in the anesthetized brain. This approach allowed us to explore how different EEG phases influence pain-related neuronal activity. Our results show major differences between the two phases of urethane-induced activity. We offer explanations for contradictory results of the previous researcher in regard to responses to painful stimuli in anesthetized animals. Furthermore, we suggest that the presented set of parameters (amplitude, relative amplitude, cross-correlation value) could be used as a simple and useful tool for EEG pain description under the state of general anesthesia.

The findings of this study have implications for both animal research and human healthcare. In the field of animal research, a better understanding of pain processing under urethane anesthesia can lead to improved experimental designs and more reliable results. For human medicine, the insights gained from this research could contribute to the development of more sophisticated monitoring techniques for assessing pain in unconscious or non-communicative patients, such as those under general anesthesia for surgical procedures or in intensive care units. Moreover, by making a step towards revealing the basic principles of pain processing under anesthesia, our results can open new avenues for neurogenic pain research and contribute to our fundamental understanding of pain physiology. The observed patterns of cortical activity in response to pain stimulation during different phases of urethane anesthesia could provide valuable clues about the underlying mechanisms of pain perception and modulation, potentially leading to novel approaches in pain management and the treatment of various pain disorders.

## Notes

### Competing Interest Statement

The authors have declared no competing interest.

